# A rat model of pregnancy in the male parabiont

**DOI:** 10.1101/2021.06.09.447686

**Authors:** Rongjia Zhang, Yuhuan Liu

**Affiliations:** Experimental Teaching Demonstration Center of Education Institutions, Faculty of Naval Medicine, Naval Medical University, Shanghai, China; Department of Nutrition and Food Hygiene, Faculty of Naval Medicine, Naval Medical University, Shanghai, China; Department of Obstetrics and Gynecology, Changhai Hospital, Naval Medical University, Shanghai, China

## Abstract

Male pregnancy is a unique phenomenon in syngnathidae which refers to embryo or fetus incubation by males. However, whether male mammalian animals have the potential to conceive and maintain pregnancy remains unclear. Here, we constructed a rat model of pregnancy in the male parabiont using a four-step strategy. First, a heterosexual parabiotic pair was produced by surgically joining a castrated male rat and a female rat. After 8 weeks, uterus transplantation was performed on the male parabiont. After recovery, blastocyst-stage embryos were transplanted to the grafted uterus of the male parabiont and the native uterus of the female parabiont. Caesarean section was performed at embryonic day 21.5. The success rate of the model was 3.68%, and pups were only delivered from the male parabiont under pregnant blood exposure. Our results reveal the potential for rat embryonic development in male parabionts, which may have a profound impact on reproductive biology.

**One-sentence summary:** A rat model of pregnancy was constructed in the male parabiont using a four-step strategy.

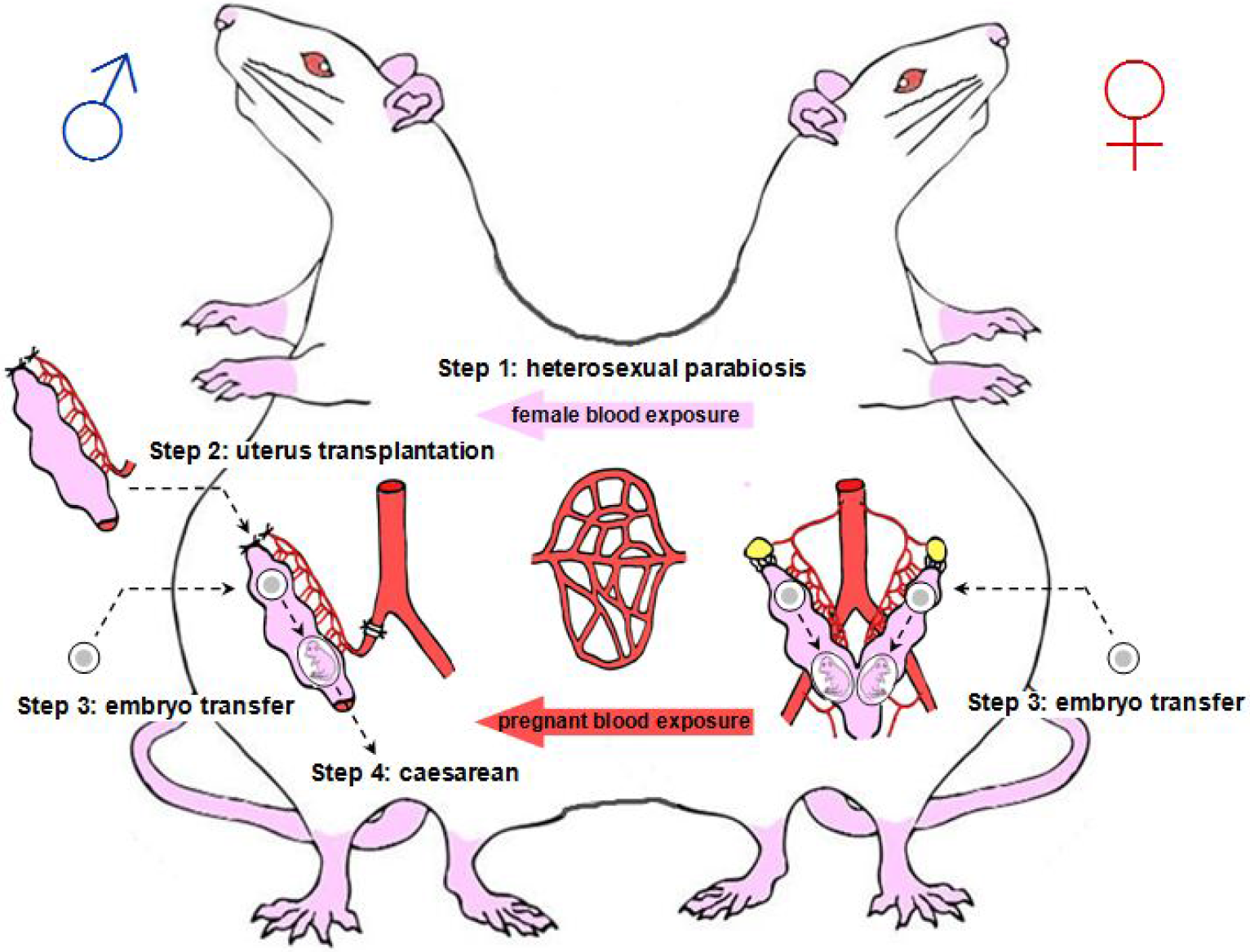

## MAIN TEXT

Male pregnancy, generally comprising embryo or fetus incubation by males until birth, is an extremely rare phenomenon in nature (*1*). Syngnathidae is the only family in which males are known to carry offspring during pregnancy (*2, 3*). In mammalian animals, pregnancies are carried by females. However, whether male mammalian animals have the potential to conceive and maintain pregnancy remains unclear. Previous studies have reported that mouse embryos transferred to non-uterine organs of male hosts develop only to a limited stage (*4, 5*), suggesting the restriction of complete embryo development in male mammalian bodies. We hypothesized that such restriction may be caused by two points: 1, lack of a uterus for embryo implantation and development; 2, lack of female and pregnant micro-environmental conditions that promote endometrial growth and allow embryo implantation or development.

In this study, we investigated whether male pregnancy followed by live birth could be achieved in a rat model if the presumptive constraints were solved using existing methods. Accordingly, uterus transplantation (UTx) and heterosexual parabiosis were conducted to test our hypothesis. UTx is a surgical procedure that has been applied in several mammalian species (*6–12*). Parabiosis is an experimental model in which two animals are surgically connected, such that their blood microenvironment is shared through anastomosis (*13, 14*). Our experiment employed a four-step strategy. First, a male rat was castrated and surgically joined with a female rat to confer a female microenvironment to the male rat through blood exchange. Next, UTx was performed on the male parabiont. Then, we transferred blastocyst-stage embryos to both the grafted uterus of the male parabiont and the native uterus of the female parabiont to observe embryonic development in the grafted uterus of the male parabiont under pregnant blood exposure. Finally, we performed caesarean section if the male parabiont was pregnant.

The first step of our experiment was to model and screen heterosexual parabiotic pairs (Fig. 1A). To reduce potential immune rejection caused by parabiosis and subsequent UTx, we selected inbred Lewis rats for use in this study. We firstly performed preliminary screening of all female and male Lewis rats. After 2 weeks of vaginal smear observation, we selected female rats who had three regular estrus-metestrus-diestrus-proestrus, estrus-metestrus-metestrus-proestrus, or estrus-diestrus-diestrus-proestrus cycles, each lasting 4 days (Fig. 1B, 1C). Female rats were excluded if they had no obvious estrous cycles or had irregular estrous cycles (Fig. 1C). The selected females were divided into two groups: donors for UTx and female parabionts for heterosexual parabiosis surgery. Male rats were mated with superovulated female rats to verify their reproductive function before selection of the male parabiont group. Prior to parabiosis surgery (Fig. S1A), the testes, epididymes, right ventral prostate, and seminal vesicles of each male were removed (Fig. 1D). Two weeks after the parabiosis surgery (Fig. 1D), both female donors and female parabionts were received estrous cycle synchronizations, and only those with three synchronized estrous cycles assessed by vaginal cytology could proceed for the next step (Fig. 1A, 1C).

**Fig. 1.**
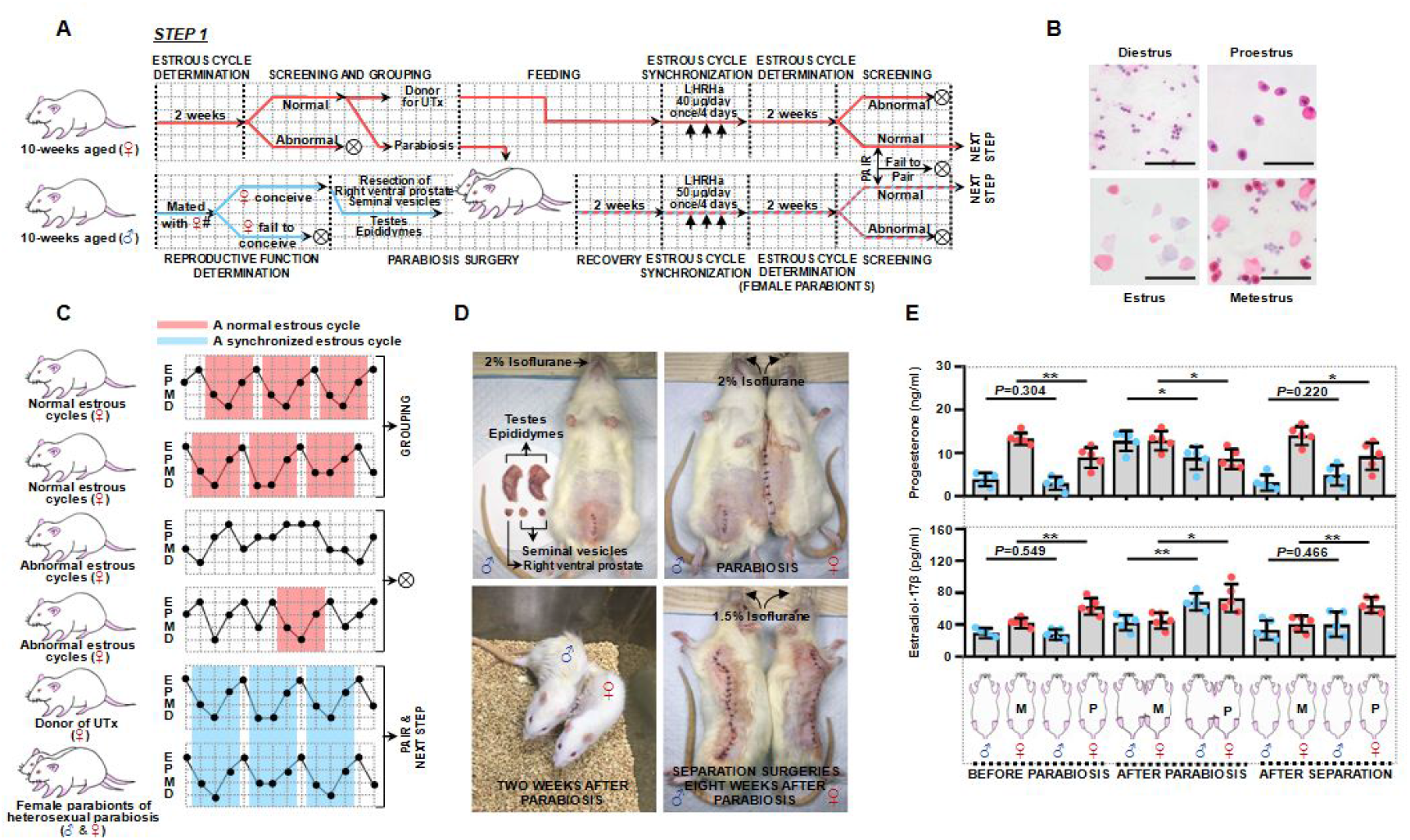
Modeling and screening of heterosexual parabiotic pairs. (A) Graph illustrating heterosexual parabiosis modeling and screening. Hashes indicate superovulated female rats. Crosses indicate rats removed from the experiment. (B) Photomicrographs of vaginal smears from female rats in diestrus, proestrus, estrus, or metestrus. Scale bars, 100 μm. (C) Representative images showing estrous cycles in female rats and female parabionts of heterosexual parabiotic pairs during a 2-week period. E, estrus; P, proestrus; M, metestrus; D, diestrus. (D) Heterosexual parabiosis and separation surgical procedures. (E) Progesterone and estradiol-17β serum levels in selected heterosexual pairs prior to parabiosis surgery, at 6 weeks after parabiosis surgery, and 2 weeks after separation surgery (n = 5 per group). M, late-stage metestrus; P, late-stage proestrus. Error bars indicate standard deviations. \**P* < 0.05; \*\**P* < 0.01.

To explore whether male parabionts were under female blood exposure following parabiosis surgery, we examined progesterone and estrogen-17β serum levels in male and female parabionts. Late-stage metestrus and late-stage proestrus were selected as time points (Fig. S1B) because they indicate hormone peaks in female rats (*15*). Female rats who exhibited three estrus-metestrus-diestrus-proestrus cycles before parabiosis surgery showed significantly different levels of both progesterone and estrogen-17β between late-stage metestrus and late-stage proestrus. However, these trends were not observed in male rats at the same time points (Fig. 1E). At 6 weeks after parabiosis surgery, female parabionts maintained significant hormone alteration, and male parabionts exhibited similar trends (Fig. 1E). To verify that the acquired hormone alterations in male parabionts were induced through exposure to blood from female parabionts, separation surgery was performed at 8 weeks after parabiosis surgery (Fig. 1D). Progesterone and estrogen-17β levels remained significantly different between late metestrus and late proestrus in female rats at 2 weeks after separation, whereas these hormone level differences between late metestrus and late proestrus were lost in male rats (Fig. 1E). We subsequently examined testosterone serum levels to determine whether female parabionts were affected by androgen from male parabionts following parabiosis surgery. At 6 weeks after parabiosis surgery, testosterone levels were significantly decreased in the male parabionts; no significant difference in testosterone levels were observed in the female parabionts (Fig. S1C).

The second step of our experiment was to transplant grafted uteruses into male parabionts within heterosexual parabiotic pairs (Fig. 2A). We devised a novel UTx protocol involving anastomosis of the right common iliac vessel of the grafted uterus with the right common lilac vessel of the male parabiont using end-to-end cuff technology (Fig. 2B) (*16–18*). However, because this surgical protocol could lead to right hindlimb ischemia, which would ultimately impair the function of the grafted uterus in the male parabiont, we first explored the effects of ligation and incision of the right common iliac artery and vein in the right hindlimb in male rats. The results showed no obvious ischemia or necrosis in the right hindlimb at 8 weeks after surgery (Fig. S2A). Additionally, we ligated and cut off the right iliac vessel of the female rats, maintaining the branch that supplies the uterus. We found no visible hindlimb ischemia or necrosis at 8 weeks after surgery (Fig. S2B). Next, we mated these female rats with normal male rats. We found that the surgical procedure did not significantly impair female fertility (Fig. S2C) and did not alter weight evolution in the resulting pups (Fig. S2D). Therefore, our UTx protocol appeared to have caused neither hindlimb ischemia nor uterine damage at 8 weeks after UTx.

**Fig. 2.**
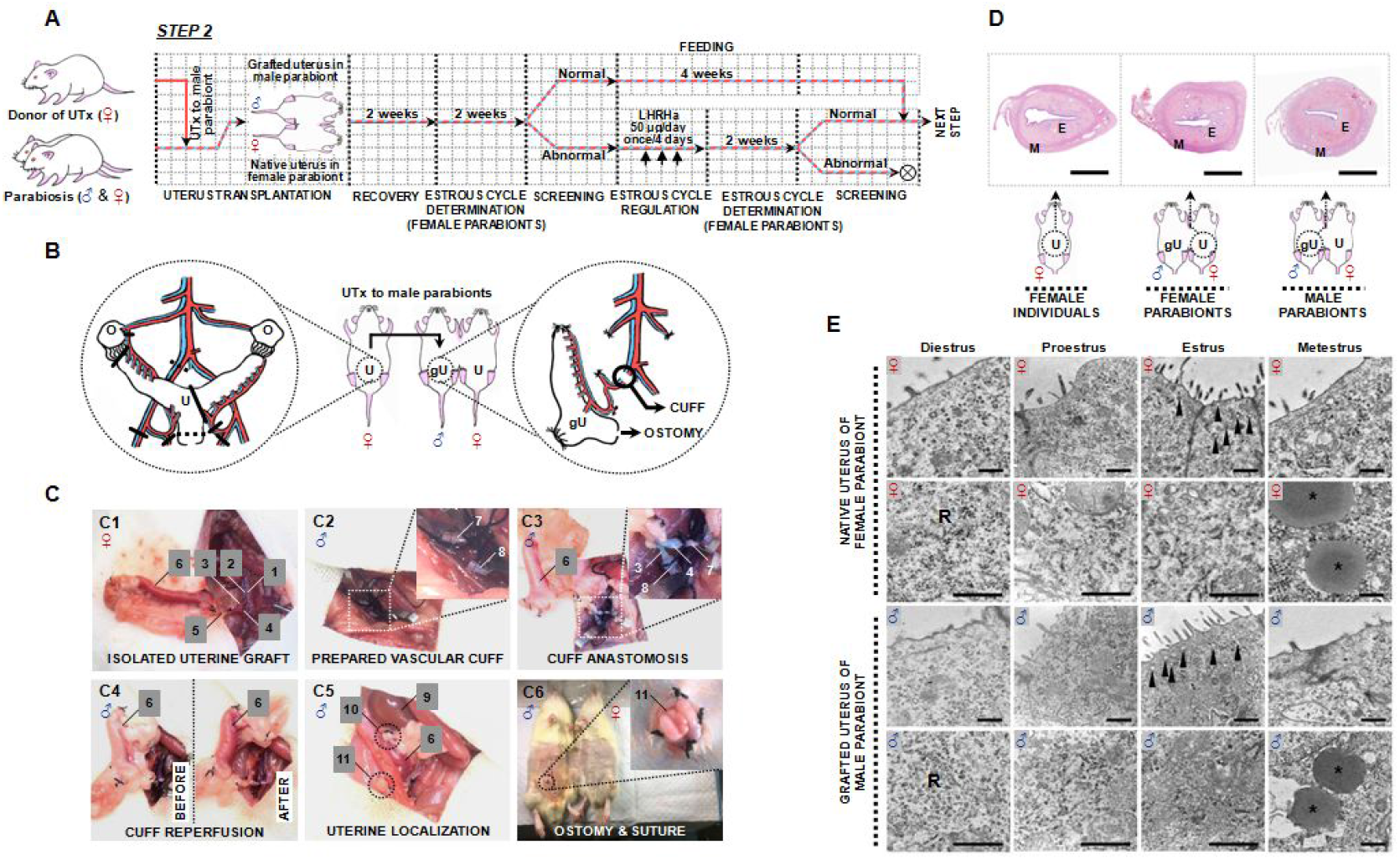
Grafted uteruses transplanted into male parabionts. (A) Illustration of uterus transplantation (UTx) and recovery. Crosses indicate rats removed from the experiment. (B) Schematic drawing of UTx. U, uterus; gU, grafted uterus; O, ovary. Solid lines indicate ligation and cuts. Dotted lines indicate cuts. (C) Surgical procedure for UTx. 1, aorta; 2, vena cava; 3, common iliac artery; 4, common iliac vein; 5, uterine vessels; 6, uterus; 7, common iliac arterial cuff; 8, common iliac venous cuff; 9, kidney; 10, connective tissue beneath kidney; 11, stoma. (D) Hematoxylin and eosin (H & E) staining images of native uteruses from female individuals, native uteruses from female parabionts, and grafted uteruses from male parabionts (8 weeks after UTx). E, endometrium; M, muscle layer. Scale bars, 1 mm. (E) Electron microscopy images of native uteruses from female parabionts and grafted uteruses from male parabionts. Diestrus, proestrus, estrus, and metestrus were determined by vaginal smears from female parabionts (n = 3 for each stage). R, free ribosomes. Arrows indicate secretory granules. Asterisks indicate lipid vacuoles. Scale bars, 500 nm.

Next, we implemented our UTx protocol based on cuff technology (Fig. S3). The uterine graft was isolated in the female donor; cuff preparation, cuff anastomosis, cuff reperfusion, uterine localization, and ostomy were then performed in the male parabiont (Fig. 2C). Prior to formal UTx surgery, we conducted surgical training to improve the UTx success rate (Fig. S4). To reduce the number of animals used in surgical training, we selected male individuals, rather than male parabionts in heterosexual parabiotic pairs, as the recipients (Fig. S4A). We divided the surgical training into two stages; we performed formal UTx after ensuring that the duration of warm ischemia–reperfusion (I/R) could be limited to ≤ 30 min (Fig. S4B–E). After formal UTx, the skin around the stoma was sutured with two gauzes and long-term care was performed (Fig. S5A, B). During recovery, we monitored the estrous cycles of the female parabionts; we provided hormonal regulation to the female parabiont if the estrous cycle was unusual. At 8 weeks after UTx, surviving heterosexual parabiotic pairs with normal estrous cycles in the female parabionts were subjected to the next step.

The functional recovery of a grafted uterus is closely related to immune rejection (*19, 20*) and I/R injury (*21*). Therefore, we performed hematoxylin and eosin (H & E) and CD8^+^ immunohistochemical staining at 8 weeks after UTx. We observed no large areas of necrosis (a typical feature of immune rejection) (*19, 20*) and no obvious blood extravasation or severe endometrial loss (typical features of I/R) (*21*) in the grafted uteruses of male parabionts (Fig. 2D). Furthermore, no significant differences in CD8^+^ counts were observed in the grafted uteruses of male parabionts, compared with the native uteruses of either female individuals or female parabionts (Fig. S5C, D). To further investigate whether the grafted uteruses of male parabionts were influenced by exposure to blood from female parabionts, we examined both grafted and native uteruses using electron microscopy at various estrous stages in female parabionts. Consistent with electron microscopy findings in a previous study (*15*), the native uteruses of female parabionts showed ultrastructures that were characteristic of the different estrous stages, and the grafted uteruses of male parabionts exhibited similar ultrastructures (Figs. 2E and S5E).

The third step of our experiment was to transplant blastocyst-stage embryos to the grafted uteruses of male parabionts and native uteruses of female parabionts (Fig. 3A). At 3 days before embryo transfer, female parabionts were mated with vasectomized male rats to expose the male parabionts to blood from pseudo-pregnant females. The vasectomized male rats selected for mating were trained and screened before vasectomy to increase the mating success rate (Fig. 3B). On the day of embryo transfer, blastocyst-stage embryos were collected from treated female rats; heterosexual parabiotic pairs were then subjected to laparotomy to observe whether the morphology and color of grafted uteruses in the male parabiont were similar to the morphology and color of native uteruses in the female parabionts (Fig. 3C). Next, embryos were transplanted to the left uteruses of the female parabionts, grafted uteruses of the male parabionts, and right uteruses of the female parabionts (Fig. S6A). Immunosuppression and stoma care were continued after embryo transfer (Fig. S5B).

**Fig. 3.**
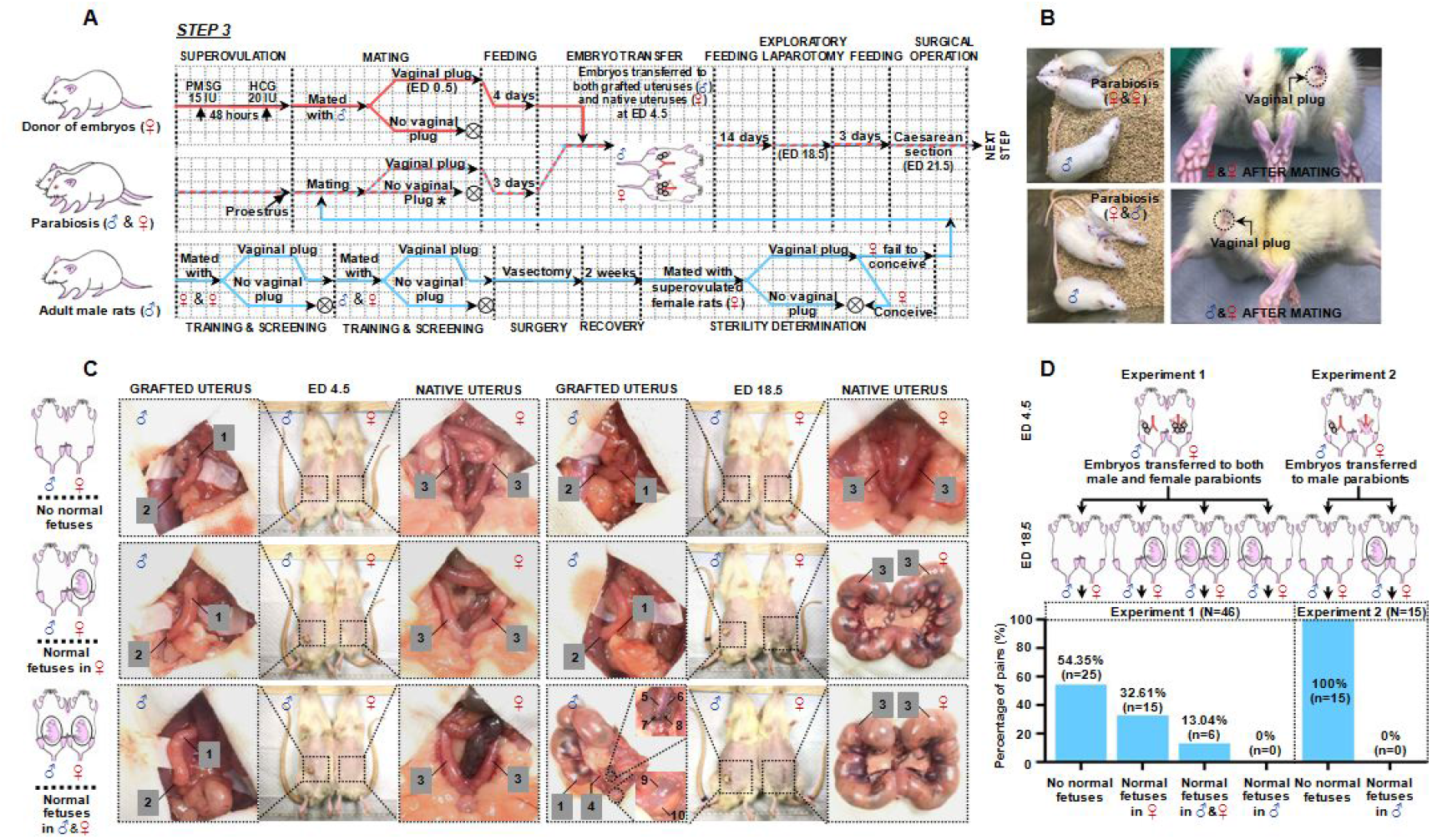
Transferred embryos developed in grafted uteruses of male parabionts during exposure to blood from pregnant female parabionts. (A) Illustration of the embryo transfer procedure. Crosses indicate rats removed from the experiment. (B) Male rats were trained for mating with female parabionts prior to vasectomy. (C) Representative heterosexual parabiotic pair, grafted male parabiont uterus, and native female parabiont uterus at embryonic day (ED) 4.5 (before embryo transfer) and ED 18.5 (exploratory laparotomy). 1, grafted uterus; 2, uterine stump close to stoma; 3, native uterus; 4, uterine vessels; 5, common iliac artery; 6, common iliac vein; 7, common iliac arterial cuff; 8, common iliac venous cuff; 9, bladder; 10, atrophied ventral prostate. (D) Pregnancy rate of heterosexual parabiotic pairs at ED 18.5. Pregnancy was defined as at least one normally developing fetus in a native uterus of the female parabiont or grafted uterus of the male parabiont.

In total, we transferred 842 blastocyst-stage embryos to 46 heterosexual parabiotic pairs at embryonic day (ED) 4.5, including 562 embryos transferred to female parabionts and 280 embryos transferred to male parabionts. At ED 18.5, exploratory laparotomy was performed to observe the development of transferred embryos (Fig. 3C). We found that 169 (30.07%) embryos had developed normally in the native uteruses of female parabionts, whereas only 27 (9.64%) embryos had developed normally in the grafted uteruses of male parabionts (Fig. S6B). Further data mining indicated that all developing embryos in the male parabionts had been exposed to the blood environment of the pregnant female parabionts (Fig. S6C). Among all heterosexual parabiotic pairs, 25 (54.35%) showed no normal embryos in either male or female parabionts (Fig. 3C and Movie 1), 15 (32.61%) showed at least one normal embryo in female parabionts alone (Fig. 3C and Movie 2), 6 (13.06%) showed at least one normal embryo in both male and female parabionts (Fig. 3C and Movie 3), and 0 showed at least one normal embryo in male parabionts alone (Fig. 3D). Therefore, we inferred that the transplanted embryos were able to develop normally in the grafted uterus of a male parabiont only during exposure to blood from a pregnant female parabiont. To test this hypothesis, we transplanted 90 blastocyst-stage embryos only to the grafted uteruses of male parabionts at ED 4.5 (n = 15). As expected, no normal developing embryos were found in the grafted uteruses of male parabionts at ED 18.5 (Figs. 3D and S6C).

The last step of our experiment was to observe the pregnant male parabionts and their surviving fetuses after caesarean section (Fig. 4A). At ED 21.5, we performed caesarean sections on pregnant female individuals who had conceived through normal copulation, which revealed survival among all fetuses. However, some of these fetuses died within 2 h (Fig. 4B), perhaps due to early termination of pregnancy. Next, we performed caesarean sections on heterosexual parabiotic pairs in which both the male and female parabionts were pregnant. Except for one resorbed fetus, all fetuses removed from female parabionts were alive; however, some died within 2 h (Fig. 4B, C). While performing caesarean section on the male parabionts, we found some abnormal dead fetuses; this phenomenon did not occur in either of the other two groups (Fig. 4B). These dead fetuses differed from normal fetuses in terms of morphology and color; they were accompanied by atrophied or swollen placentas. Caesarean section of the male parabionts also produced a small number of resorbed fetuses and surviving fetuses that survived for 2 h (Fig. 4B) and 24 h (Movie 4). At 2 h after caesarean section, the body weight and placental weight of live fetuses removed from male parabionts did not significantly differ from the body weight and placental weight of live fetuses in the other two groups (Fig. 4C). These newborn pups from male parabionts developed normally to adulthood and were functionally reproductive (Figs. S7A, B, and 4E). Histological examination showed no obvious abnormalities in the heart, lung, liver, kidney, brain, testis, epididymis, ovary, or uterus of offspring successfully delivered from male parabionts by caesarean section (Figs. S7C and 4D).

**Fig. 4.**
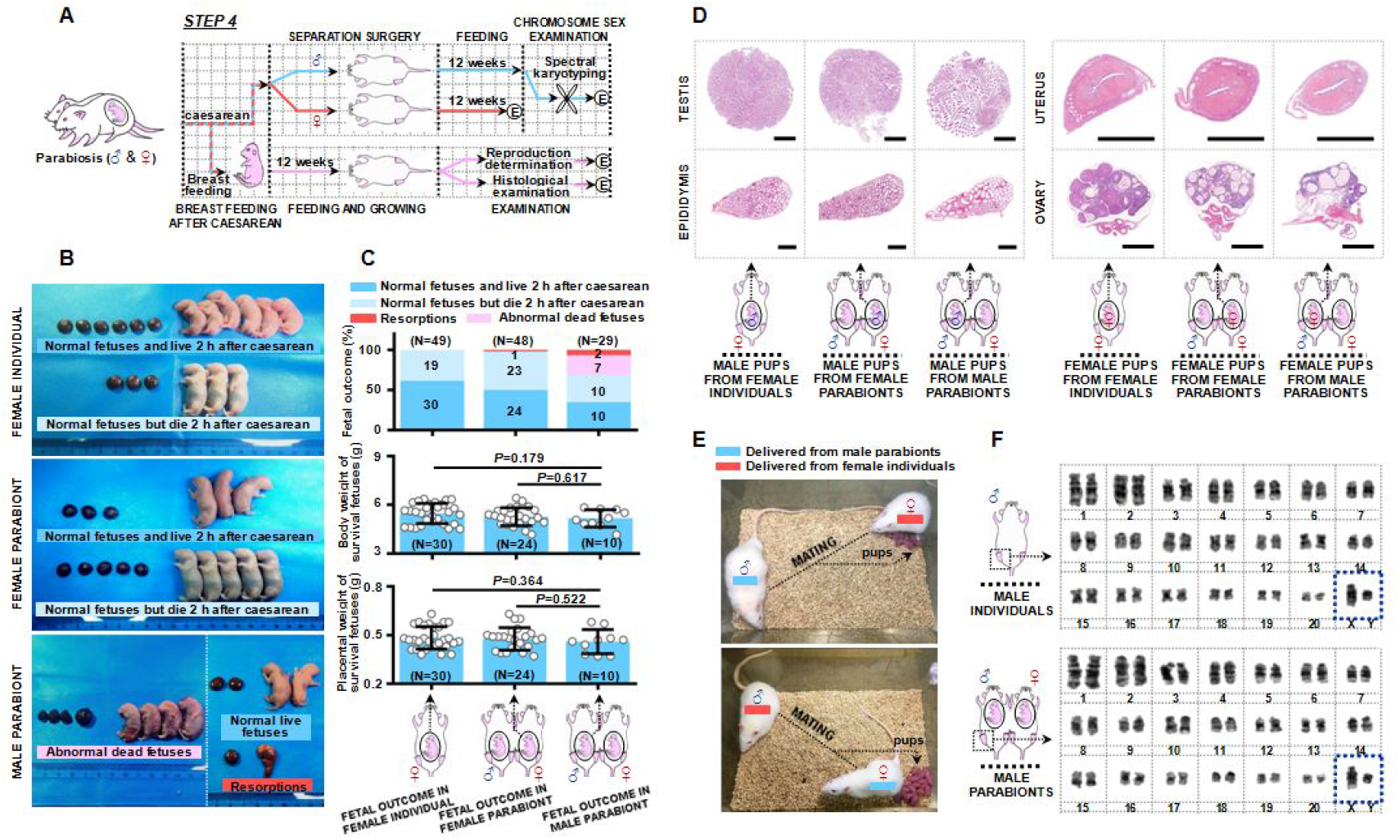
Surviving fetuses and male parabionts after caesarean section. (A) Illustration of the experimental protocol following caesarean section. E, euthanization. (B) Representative images of fetuses and placentas at 2 h after caesarean section delivery from female individuals, female parabionts, and male parabionts. (C) Fetal outcomes following caesarean section. The body weight and placental weight of surviving normal fetuses at 2 h after caesarean section are shown. Error bars indicate standard deviations. (D) H & E staining images of the testis and epididymis of male offspring (12 weeks; scale bars, 5 mm) and the uterus and ovary of female offspring (12 weeks; scale bars, 2 mm). (E) Male and female fetuses delivered from male parabionts developed to adulthood and exhibited reproductive competence. (F) Representative images of spectral karyotyping (femoral bone marrow). In this analysis, male individuals were defined as male rats who showed reproductive competence prior to castration and received castration surgery at 3 months of age; male parabionts were defined as rats who had experienced successful pregnancy and survived following separation surgery. Both male individuals and male parabionts were sampled for karyotype analysis at 12 months of age.

After caesarean section, we performed separation surgery on the heterosexual parabiotic pairs. All separated male parabionts survived for at least 3 months after surgery. We performed karyotype analysis on the separated male parabionts to determine their chromosomal sex, using normal male individuals (no treatment after castration) as a reference. Although karyotype results can change in inbred rats (*22*), we found no significant difference in sex chromosomes between separated male parabionts and normal male individuals (Fig. 4F), suggesting that the chromosomal sex of the separated male parabionts was indeed male.

To our knowledge, pregnancy has never been reported in male mammalian animals. In this study, we constructed a rat model of pregnancy in the male parabiont and found that transplanted blastocyst-stage embryos were able to develop to adulthood in the grafted uteruses of male parabionts during exposure to the blood of pregnant female parabionts. The success rate of the entire experiment was very low, but 10 pups were successfully delivered from male parabionts by caesarean section and developed to adulthood (Fig. S8). We also discovered three new phenomena in our rat model. First, during caesarean section at ED 21.5, abnormal dead fetuses were observed in the grafted uteruses of male parabionts. Because no abnormalities were found during laparotomy at ED 18.5, we infer that these abnormal fetal deaths began during late embryonic development (ED 18.5–21.5) in these male parabionts. It remains unknown whether this phenomenon is peculiar to pregnancy in male parabionts. Second, embryos might develop normally in male parabionts only when exposed to blood from pregnant female parabionts, not to blood from non-pregnant females. The specific mechanism requires further investigation. Third, we noticed a detail that female parabionts might exhibit non-pear shape enlarging during pregnancy. This may be due to the squeezing of the space occupied by the male parabionts, or exposure of castrated male blood adversely affects the signs of female parabionts. It seems that castrated male parabionts and female parabionts can influence each other by blood exchange, and this is why female parabionts may still lose their normal estrous cycles after parabiosis surgery. Their relationship needs further research.

Finally, our findings reveal the potential for rat embryonic development in male parabionts, and it may have a profound impact on reproductive biology research.

## Acknowledgments

We are grateful to Jianan Tang (Fudan University) and Fangxing Lin, Yaping Zhou, Yiqing Zhu (Naval Medical University) for technical assistance and to Ling Han, Hongtao Lu, Hui Shen, Su Liu (Naval Medical University) for administrative assistance. We thank Xuejun Sun (Naval Medical University) for constant encouragement and help. This work was supported by the National Natural Science Foundation of China (Grant No. 81500480 and No. 81641059).

## Author contributions

R.Z. conceived the project. R.Z. designed and performed the experiments with the aid of Y.L.. R.Z. and Y.L. analyzed the data and wrote the manuscript.

## Data and material availability

All data are available in the main text or in the supplementary materials.

## Materials and Methods

### Animal Ethics

All experimental procedures were performed according to local ethics guidelines on animal welfare and permits which can be summarized as reducing the number of animals used and minimizing animal suffering. Thus, we selected male individuals as the recipients in surgical training, and the number of animals used has been reduced by 60. All manipulations were performed under anesthesia and none of the rats exhibited signs of pain during the course of the experiment. All surgical protocols were designed according to previous reports (*4–7, 13, 14, 16–21, 27*), and caesarean section was performed to avoid labor pain.

### Animal housing and parabiosis care

All male and female inbred Lewis rats were obtained from Shanghai Sippr-BK Laboratory Animal Co., Ltd. and housed in the animal experimental center of the Naval Medical University and the experimental teaching center of the Facility of Naval Medicine. The animals were maintained under a 12-h/12-h light/dark cycle with free access to food and water. Parabiotic pairs were housed individually to avoid biting. To reduce wound infection induced by faecalis adherence, we replaced the padding daily within the first 3 days after surgery and at 3-day intervals for the remainder of the experiment.

### Vaginal cytology for estrous cycle determination

The estrous cycles of female rats were monitored for 14 days using vaginal cytology. Vaginal cytology was examined once daily at approximately 8:00 a.m. by collection of vaginal secretions using sterile plastic pipettes filled with saline. Vaginal fluid was placed onto glass slides and fixed by adding a drop of 95% alcohol. The slides were then stained with H & E and examined under a microscope (Olympus, CX23). Female rats were considered to be in diestrus, proestrus, estrus, or metestrus based on the following proportions of cell types: a diestrus smear consisted primarily of leukocytes; a proestrus smear consisted primarily of nucleated epithelial cells; an estrous smear consisted primarily of nucleated cornified cells; and a metestrus smear consisted of equal proportions of leukocytes, cornified epithelial, and nucleated epithelial cells (*23*).

### Heterosexual parabiosis and separation surgery

For heterosexual parabiosis surgery, both male and female rats were anesthetized with isoflurane (5% for induction, 2% for maintenance, 2 L/min oxygen) (*24*) and placed in the supine position. Laparotomy was performed on the male rat through midline incision. The bilateral testes and epididymes of male rats were removed to reduce the effects of androgens on female parabionts. Resection of the seminal vesicles was performed to avoid false positive results from the vaginal plug during the mating of female parabionts and sterilized male rats (Fig. 3A). The right ventral prostate was removed to avoid physical contact with the grafted uterus. Skin incisions were made along the lateral surface in both male and female rats. Each parabiotic pair was generated by suturing the opposing flanks of the elbows and knees and then stapling the dorsal and ventral skin (Fig. S1A). After surgery, both the male and female parabionts were treated with ampicillin (20 mg/kg) to prevent infection. Isosexual (female–female) parabiotic pairs were prepared in the same manner.

For separation surgery, heterosexual parabiotic pairs were anesthetized with isoflurane (5% for induction, 1.5% for maintenance, 2 L/min oxygen). All remaining sutures were removed, and the connected skin and joints were carefully separated. Body temperature was controlled using a thermostatically heated pad.

### Blood collection for enzyme-linked immunosorbent assay

Blood samples were collected in late metestrus and late proestrus from female rats or female parabionts who had three estrus-metestrus-diestrus-proestrus cycles. Blood samples were collected from male rats at the same time points. To estimate the approximate timing of late metestrus and late proestrus, we recorded the female estrous cycles according to vaginal smear results obtained over a 14-day period (Fig. S1B). Vaginal cytology was examined at 8:00 a.m. and 4:00 p.m. on the estimated day and at 8:00 a.m. on the following day. If the females were in metestrus or proestrus at both 8:00 a.m. and 4:00 p.m. on the estimated day, we collected blood samples from the orbital venous plexus at approximately 4:30 p.m. If the females were in diestrus or estrus at 8:00 a.m. on the next day, we used blood samples collected on the previous day. Blood progesterone, estrogen-17β, and testosterone levels were measured using enzyme-linked immunosorbent assay kits (progesterone, Biovision, K7416-100; estrogen-17β, Sigma, ab108667; testosterone, Cayman, 582701), in accordance with the manufacturers' instructions. Absorbance was measured at 405 nm with correction at 600 nm.

### UTx protocol

#### Donor preparation and operation

Female donors received a luteinizing hormone-releasing hormone agonist (*25*) for estrus synchronization (Fig. 1A). UTx was performed when the female donor was in diestrus or metestrus. The donor was anesthetized with isoflurane and midline laparotomy was performed. The right uterine horn with the right uterine cavity was separated, along with a vascular pedicle including the right uterine and common iliac vessels (Fig. 2B, 2C1). After the separated tissues were confirmed to exhibit no bleeding points, a piece of warmed (37°C) saline-soaked sterile gauze was used to cover the incision.

#### Recipient preparation and operation

Female parabionts received luteinizing hormone-releasing hormone agonist for estrus synchronization (Fig. 1A). UTx was performed on the male parabiont when the female parabiont was in diestrus or metestrus. The parabiotic pair was anesthetized with isoflurane and a midline incision was made in the male parabiont. Following exposure of the right common iliac artery, the proximal portion was clipped using a microvascular clamp, the distal portion was ligated with 4-0 silk thread, and the right common iliac artery was cut between the ligature and the clip. The proximal end was flushed with sodium heparin saline (200 IU/mL); the right common iliac artery was passed through a well-designed cuff (Fig. S3A, B). The artery was then folded over the cuff and secured with an 8-0 nylon tie to expose the endothelial surface under a dissecting microscope (Hotry, MT-1). The right common iliac vein of the male parabiont was treated in a similar manner (Fig. 2C2).

#### Vascular anastomosis, vaginal ostomy, immunosuppression, and stoma care

In female donors, the right common iliac artery and vein were cut. The uterine graft was harvested and flushed with ice-cold heparin sodium saline (200 IU/mL) containing dissolved hydrogen (1.8 mg/L) to reduce I/R injury (*26*). A dissecting microscope was used to place the right common iliac artery of the graft over the arterial cuff of the male parabiont and fix it with 5-0 silk thread. The same procedure was performed to fix the right common iliac vein on the venous cuff (Figs. 2C3 and S3C). After unclamping, blood supply was restored to the uterine graft (Fig. 2C4). Adipose tissue near the uterine horn was then fixed onto the connective tissue beneath the right kidney (Fig. 2C5), and the vaginal end was everted to form a stoma (Fig. 2C6). The midline incision was closed; the skin around the stoma was sutured and fixed with sterile gauze and zinc oxide gauze (Fig. S5A). Both male and female parabionts were treated with ampicillin (20 mg/kg) and tacrolimus (0.2 mg/kg, Abcam, ab120223) (*20*). Stoma care (dressing change) and immunosuppression (0.1 mg/kg tacrolimus) were maintained throughout the remainder of the experiment, in accordance with our protocol (Fig. S5B).

### Hindlimb ischemia

Experiments were performed on adult male rats and virgin female rats with normal estrous cycles. Isoflurane anesthesia was induced in male rats; the right common iliac artery and vein were then ligated below the bifurcation from the abdominal vessels. Observations were performed 8 weeks after surgery. Isoflurane anesthesia was induced in female rats; the right common iliac artery and vein were then ligated below the uterine artery and vein. Observations were performed 8 weeks after surgery. followed by superovulation. For mating, each superovulated female rat was provided two opportunities to mate with a male rat who had demonstrated reproductive success.

### Histology and transmission electron microscopy

Anesthetized animals were perfused transcardially with saline, followed by 4% paraformaldehyde. The uterus was removed and fixed in 4% paraformaldehyde. For histological evaluation, the tissue was dehydrated, infiltrated, embedded, sectioned (4 μm), and stained with H & E or an anti-CD8 antibody (Abcam, ab237709). Sections were examined in a blinded manner, using a computer image analysis system (Kongfoong, KF-PRO-120). Transmission electron microscopy was performed on heterosexual parabiotic pairs in which the female parabionts had three estrus-metestrus-diestrus-proestrus cycles, as determined by vaginal cytology. Briefly, the tissue was post-fixed in 1% osmium tetroxide, dehydrated, infiltrated, embedded, trimmed, sectioned (70 nm), and stained with acetate double oxygenic uranium and lead citrate. Image acquisition and analysis were performed in a blinded manner, using a transmission electron microscope (Hitachi, H7650).

### Embryo transfer

#### Collection of blastocyst-cell stage embryos

Adult female Lewis rats were stimulated to undergo superovulation through intraperitoneal injection of 15 IU pregnant mare serum gonadotropin (RP17827210000, BioVendor R & D); 48 h later, they underwent intraperitoneal injection of 20 IU human chorionic gonadotropin (9002-61-3, Sigma) (*27*). The superovulated female rat and an adult male rat were allowed to mate; females who showed vaginal plugs the next morning were recorded as ED 0.5 and kept for subsequent embryo collection. At ED 4.5, the uterus was excised from the mated female rat and flushed with M2 medium (M7167, Sigma). The flushed blastocyst-cell stage embryos were selected under a dissecting microscope and rechecked by an inverted microscope (BM, BM-37XC). Then they were transferred into 200- of M2 medium supplemented with 0.5 mg/mL hyaluronidase (H4272, Sigma) for 3 min, washed with fresh M2 medium, transferred into 400-μL drops of M16 medium (M7292, Sigma), and kept in a CO_2_ incubator (37°C, 5% CO_2_, 95% air) for embryo transfer.

#### Preparation of vasectomized male rats

To increase the mating success rate, adult male rats were first mated with female homosexual parabiotic pairs and then mated with heterosexual parabiotic pairs that had not undergone UTx. Male rats who had successfully impregnated the female parabionts of both homosexual and heterosexual parabiotic pairs (i.e., vaginal plugs observed) were subjected to vasectomy (Fig. 3B). The sterility of vasectomized males was confirmed 2 weeks later (Fig. 3A).

#### Embryos transferred to heterosexual parabiotic pairs

Approximately 8 weeks after UTx, surviving heterosexual parabiotic pairs with the female parabiont in proestrus were selected and caged with a trained sterile male rat. On the following morning, female parabionts with vaginal plugs were selected for embryo transfer. Three days later, blastocyst-cell stage embryos were transferred to both the grafted uteruses of male parabionts and native uteruses of female parabionts. Briefly, the heterosexual parabiotic pairs were anesthetized with isoflurane, and frontal incisions were made in the abdominal skin of both rats in each pair. The adhesive tissues were separated; the native uterus of the female parabiont and the grafted uterus of male parabiont were then fully exposed. Under a dissecting microscope, small holes were made in the native and grafted uteruses using a needle. Then, 6–10 blastocyst-cell embryos were transplanted into the left uterus of the female parabiont, grafted uterus of the male parabiont, and right uterus of the female parabiont, respectively (Fig. S6A). Finally, the body wall and skin were sewn. A thermostatically heated pad was used to maintain body temperature during and after surgery until full recovery from anesthesia.

#### Exploratory laparotomy after embryo transfer

Heterosexual parabiotic pairs were subjected to exploratory laparotomy at 14 days after embryo transfer (ED 18.5). Under anesthesia, the abdomens of the female and male parabionts were incised, adhesive tissues were carefully isolated, and both native and grafted uteruses were exposed. Normally developing embryos and absorbed embryos were counted, and the abdominal walls were sutured.

### Caesarean and breastfeeding

Caesarean section was performed on those heterosexual parabiotic pairs which both male and female parabionts were pregnant at ED 21.5. Under anaesthesia, the native uterus of the female parabiont was removed after ligation and incision of the bilateral uterine vessels and ovarian vessels. Fetuses were removed from the uterus and placed in a 37°C saline bath on a heated pad (29–31°C) for 2 h. Concurrently, the grafted uterus and stoma of the male parabiont were removed after ligation and incision of the right common iliac vessels above the cuff. The placentas were also isolated from the uterus and maintained in a manner similar to the approach used for fetuses. The abdominal walls were closed, followed by separation surgery and antibiotic therapy (20 mg/kg ampicillin). After 2 h, the fetuses were counted and their statuses were evaluated; all fetuses and placentas were weighed. Live fetuses were then fostered by surrogate mothers who had given birth to healthy litters within 24 h.

### Spectral karyotyping

The rats were anesthetized 3 h after treatment with 100 μg/kg colchicine. After exposure and removal of the femoral joint heads, bone marrow cells were collected by washing with 0.85% saline. The suspension was subjected to hypotonic treatment with 0.075 mol/L KCL at 37°C, fixed in methanol–glacial acetic acid (3:1), and dropped onto clean wet slides. The chromosomes were dyed with Giemsa solution (no. 329757421, Sigma) and their morphological characteristics were observed and analyzed using an automatic scanning microscope and image analysis system (GSL-120, Leica) with minor artificial revisions (*22*).

### Statistical and image analyses

Statistical analyses were performed using PASW Statistics software, version 18.0; graphs were generated using GraphPad Prism software, version 8.0. Student's *t*-test was used to compare measurement data between two groups. The numbers of CD8^+^ cells were estimated in five random fields for each animal and analyzed using the Wilcoxon test. The chi-squared test or Fisher's exact test was used to compare proportions. Data are presented as means ± standard deviations. Statistical significance was evaluated at a level of *P* < 0.05.

## Supplementary Materials

**Movie 1.** A non-pregnant female parabiont (red mark) and non-pregnant male parabiont (blue mark) at embryonic day (ED) 18.5.

**Movie 2.** A pregnant female parabiont (red mark) and non-pregnant male parabiont (blue mark) at ED 18.5.

**Movie 3.** A pregnant female parabiont (red mark) and pregnant male parabiont (blue mark) at ED 18.5.

**Movie 4.** Surviving fetuses delivered from male parabionts at 24 h after caesarean section.

**Fig. S1.**
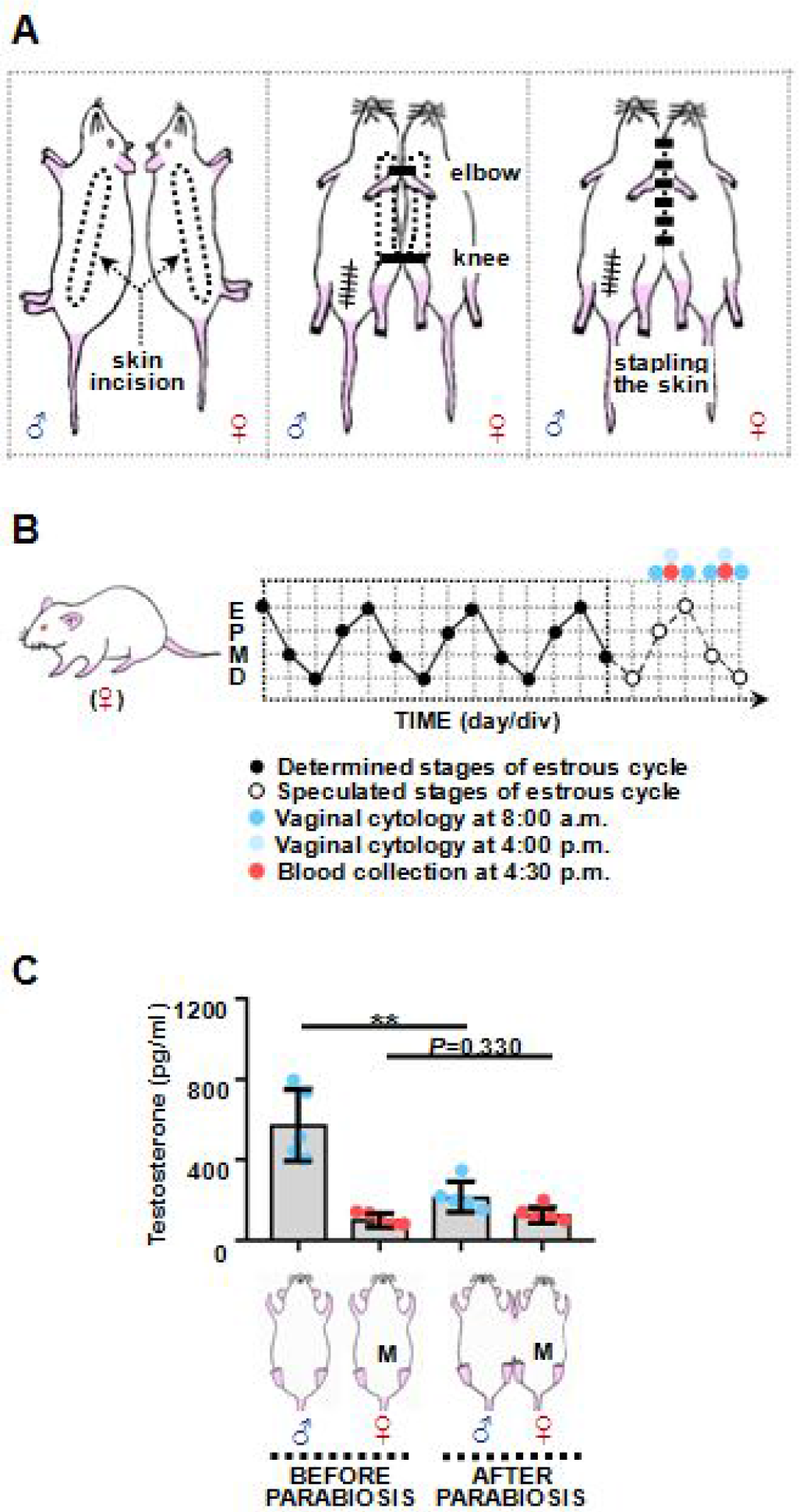
Parabiosis surgery, blood collection, and testosterone serum levels. (A) Illustration of heterosexual parabiosis surgery. After resection of the testes, epididymes, seminal vesicles, and right ventral prostate, a skin incision was made along the opposing flanks. A connection was then secured by suturing the elbows and knees using a suture pass through the soft tissue of each joint. Finally, the ventral and dorsal skin were stapled. (B) Illustration of blood collection during late-stage metestrus and late-stage proestrus. The timing of these stages was estimated by recording the female estrous cycles based on vaginal smear results obtained over a 14-day period. E, estrus; P, proestrus; M, metestrus; D, diestrus. (C) Testosterone serum levels in selected heterosexual pairs before and after parabiosis surgery (n = 5 per group). M, late-stage metestrus. Error bars indicate standard deviations. \*\**P* < 0.01.

**Fig. S2.**
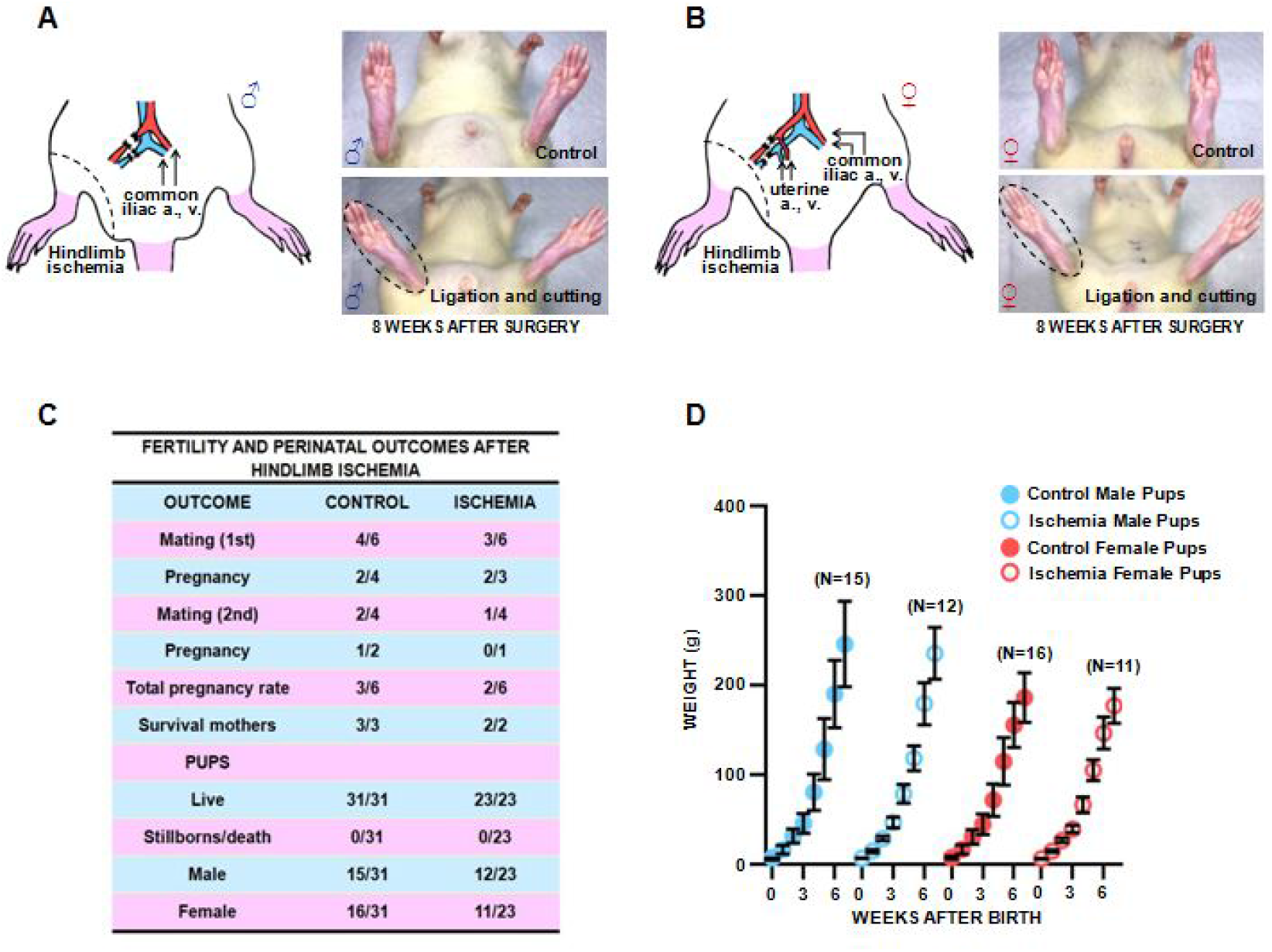
Effect of right common iliac vessel ligation and incision on hindlimb ischemia and female fertility in Lewis rats. (A and B) Schematic diagram illustrating ligation and incision of right common iliac vessels in male and female rats. Right hindlimb (dotted circle) was observed at 8 weeks after surgery. Controls were not subjected to any surgical manipulation. (C) Female fertility and perinatal outcomes after ligation and incision of right common iliac vessels. Mating was defined as successful mating, determined by visible vaginal plugs; females who were not impregnated in the first round were provided a second opportunity for mating. (D) Weight evolution of pups from different experimental groups.

**Fig. S3.**
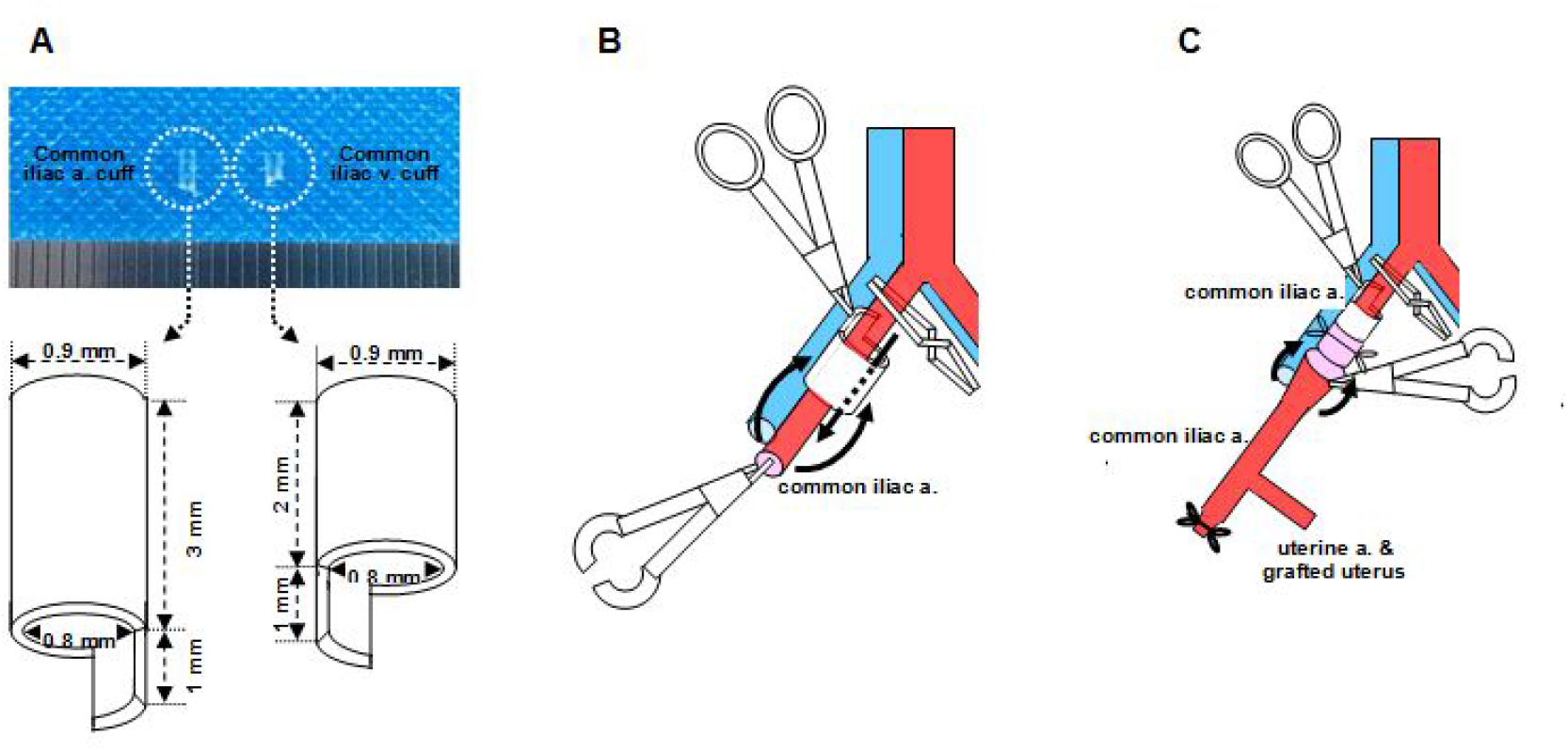
Cuff technique for establishing vascular anastomosis. (A) Vascular cuff lengths and thicknesses. (B) In the cuff technique, the common iliac artery of the male parabiont was initially pulled through the cuff. The artery was then folded over the cuff and secured with an 8-0 nylon tie to expose the endothelial surface. The common iliac vein of the male parabiont was treated in the same manner. (C) The cuffed common iliac artery was inserted into the donor common iliac artery and secured with another 8-0 nylon tie, then re-secured with a 5-0 suture. The cuffed common iliac vein of the male parabiont was treated in the same manner.

**Fig. S4.**
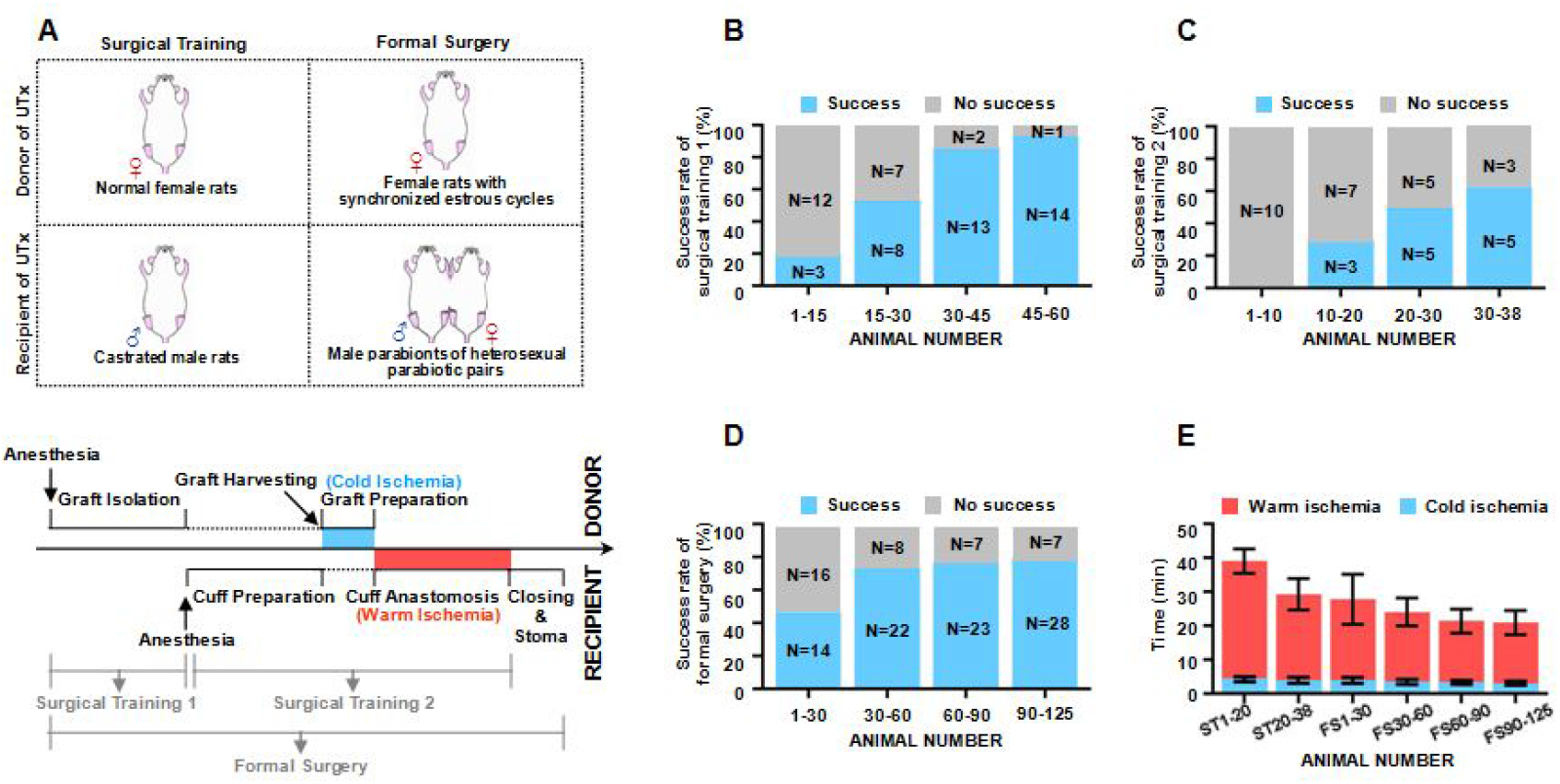
Success rate and graft ischemia duration in surgical training and formal UTx surgery. (A) Illustration of surgical training 1, surgical training 2, and formal UTx surgery. Surgical training and formal UTx surgery were performed as shown in Figs. 2B and S3, but with different donors and recipients. Success rates of (B) surgical training 1, (C) surgical training 2, and (D) formal UTx surgery. (E) Graft ischemia duration in surgical training 2 and formal UTx surgery. ST, surgical training; FS, formal UTx surgery.

**Fig. S5.**
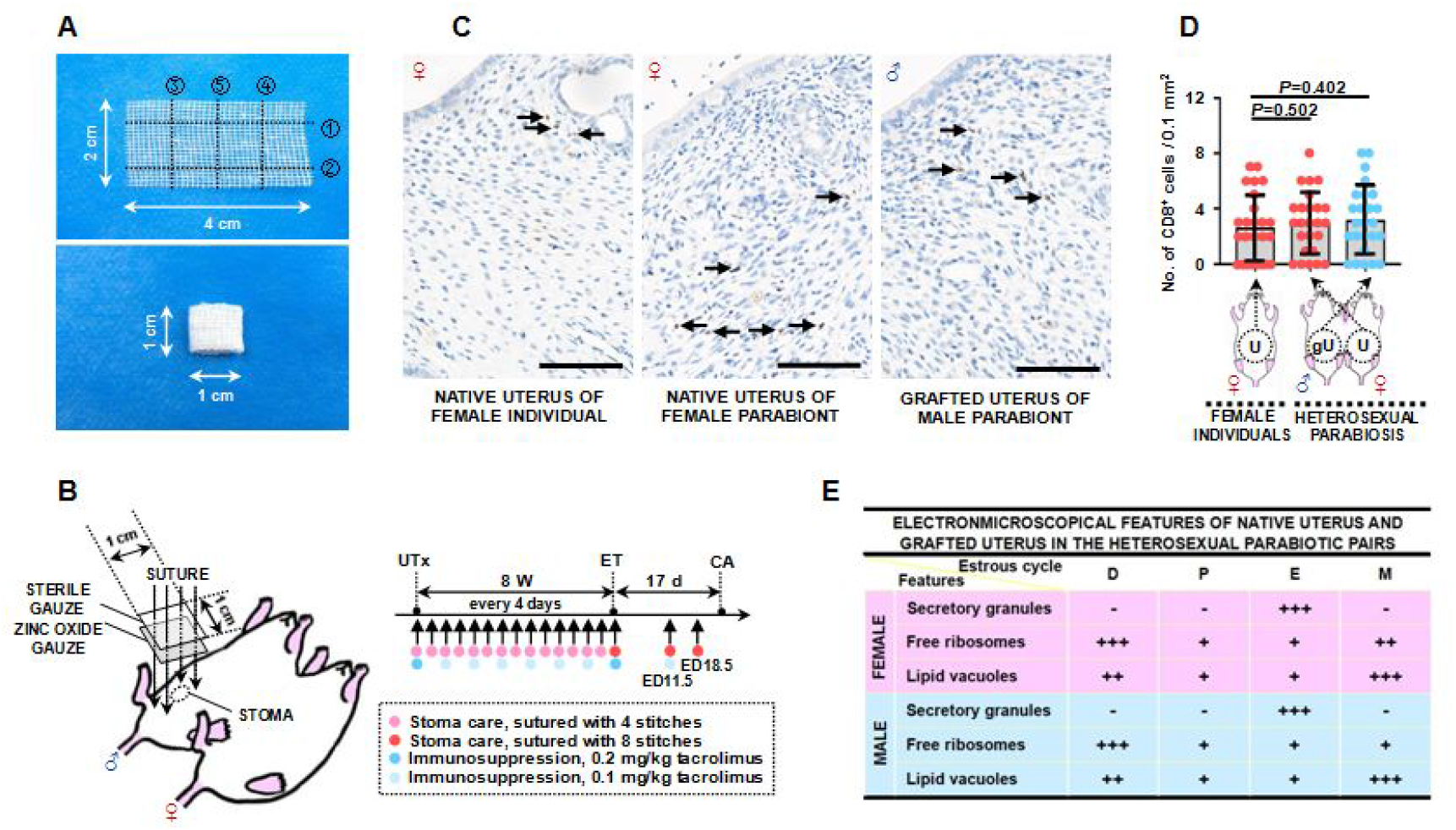
Stoma care, immunosuppression, and uterine electron microscopy features following UTx. (A) Rectangular gauze (4 cm × 2 cm) was folded into a square (1 cm × 1 cm; black dotted line). (B) Protocols for stoma care (dressing change) and immunosuppression (tacrolimus injection) from UTx until caesarean section. ET, embryo transfer; CA, caesarean section. (C) CD8 immunohistochemical staining images of native uteruses from female individuals, native uteruses from female parabionts, and grafted uteruses from male parabionts at 8 weeks after UTx. Arrows indicate CD8^+^ cells. Scale bars, 100 μm. (D) Density of CD8^+^ cells (n = 5 per group). Each rat was estimated in five random fields. U, uterus; gU, grafted uterus. (E) Major features of native uteruses from female parabionts and grafted uteruses from male parabionts observed by electron microscopy at 8 weeks after UTx (n = 3 for each estrous stage). D, diestrus; P, proestrus; E, estrus; M, metestrus. Stages were determined from vaginal smears of female parabionts. + indicates relative frequency of a feature; – indicates its total absence.

**Fig. S6.**
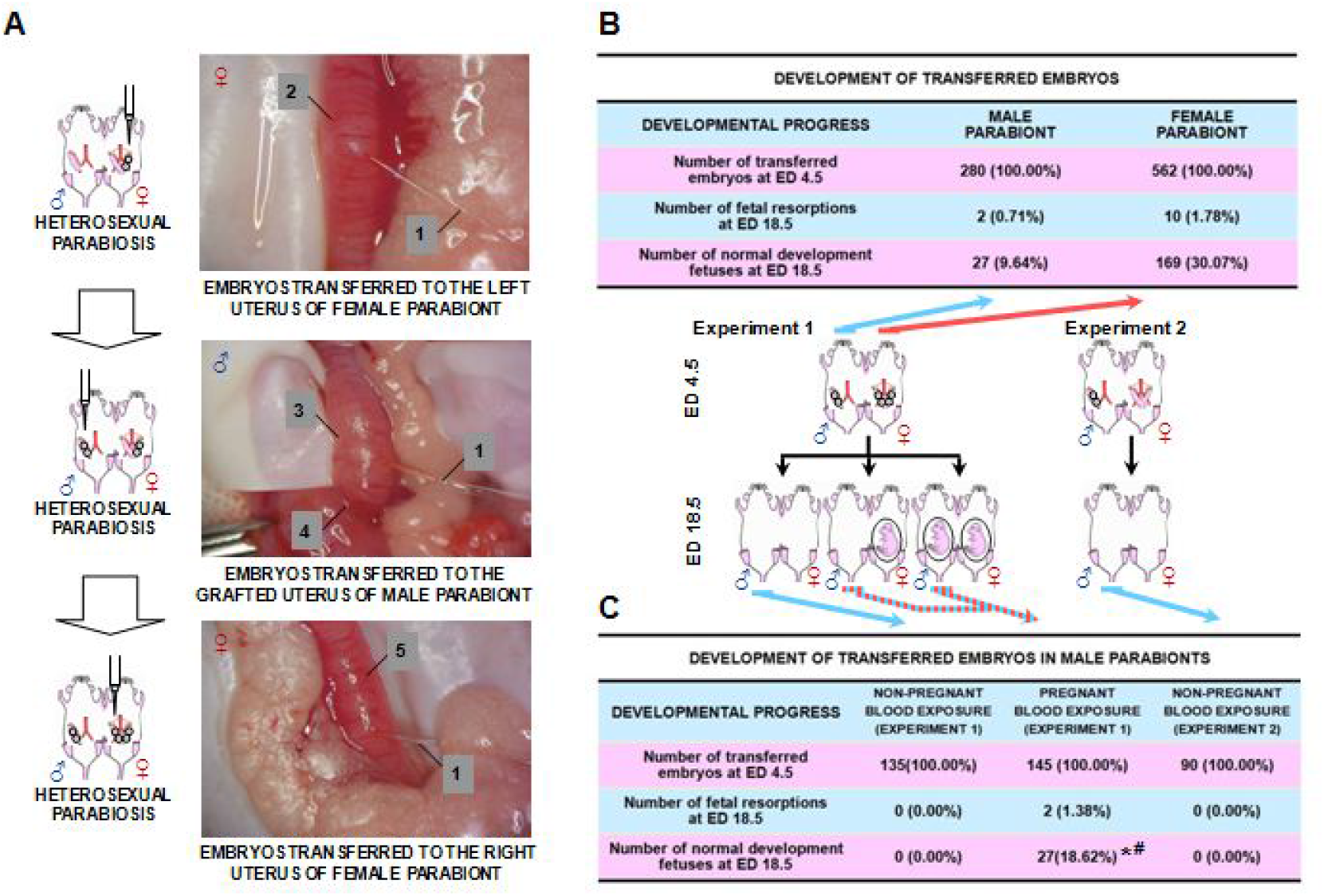
Development of transferred embryos at ED 18.5. (A) At ED 4.5, embryos were transplanted into the left uteruses of female parabionts, grafted uteruses of male parabionts, and right uteruses of female parabionts, respectively. 1, embryo pipette; 2, left uterus of female parabiont; 3, grafted uterus of male parabiont; 4, uterine stump close to stoma; 5, right uterus of female parabiont. (B) Numbers of transferred embryos at ED 4.5, fetal resorptions at ED 18.5, and normally developed fetuses at ED 18.5 in male and female parabionts. In Experiment 1, embryos were transferred to both male and female parabionts at ED 4.5; in Experiment 2, embryos were transferred only to male parabionts at ED 4.5. (C) Effects of exposure to blood from pregnant females (Experiment 1) or non-pregnant females (Experiments 1 and 2) on the development of embryos transferred to male parabionts. \**P* < 0.001 compared with exposure to blood from non-pregnant females (Experiment 1); #*P* < 0.001 compared with exposure to blood from non-pregnant females (Experiment 2).

**Fig. S7.**
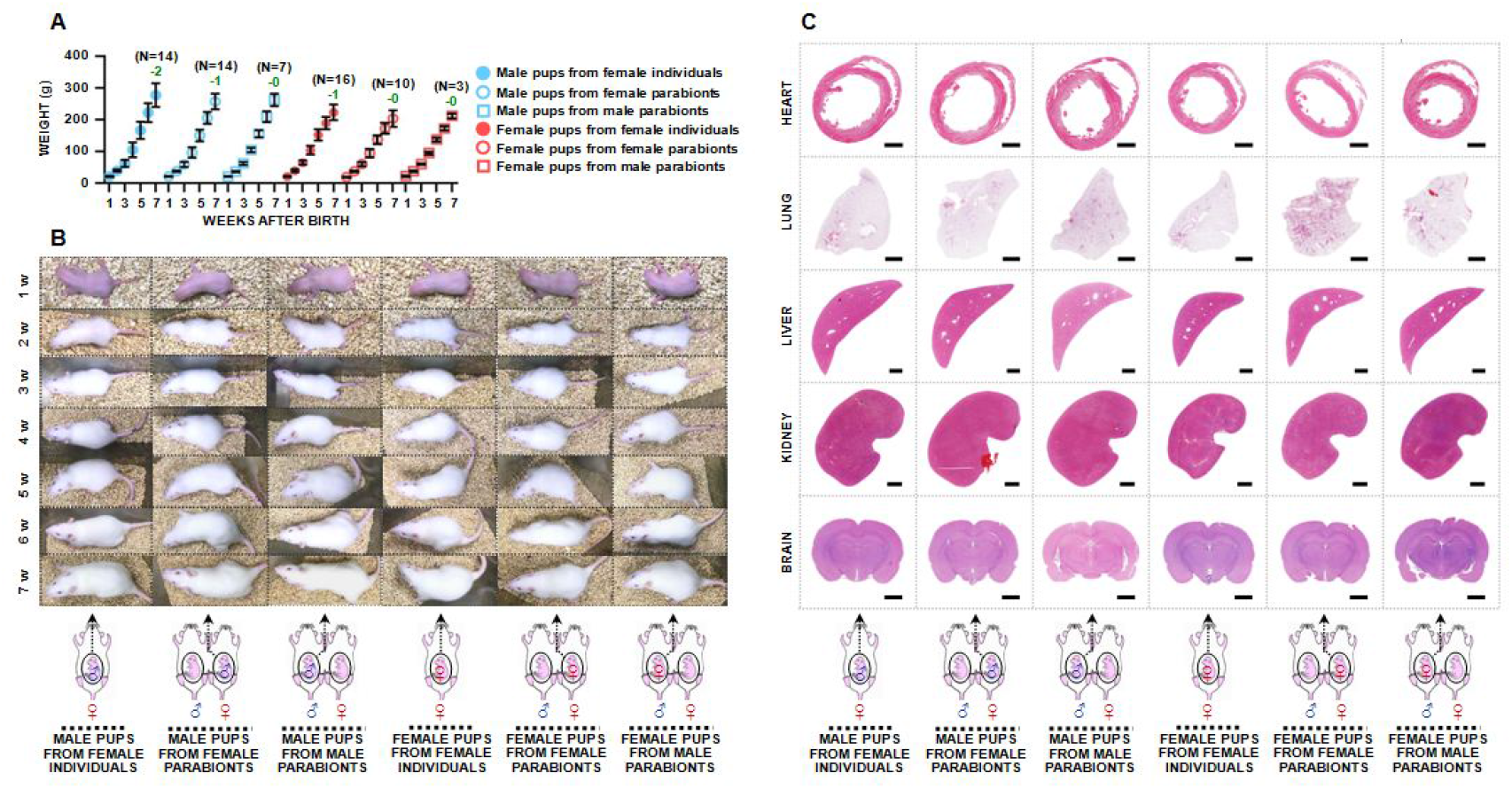
Growth of surviving fetuses after caesarean section. (A) Weight evolution of survival fetuses. Green numbers indicate numbers of pups eaten by foster mothers during breastfeeding. (B) General observations of surviving fetuses during growth and development. (C) H & E staining images of hearts, lungs, livers, kidneys and brains of male and female offspring at 12 weeks. Scale bars, 2 mm.

**Fig. S8.**
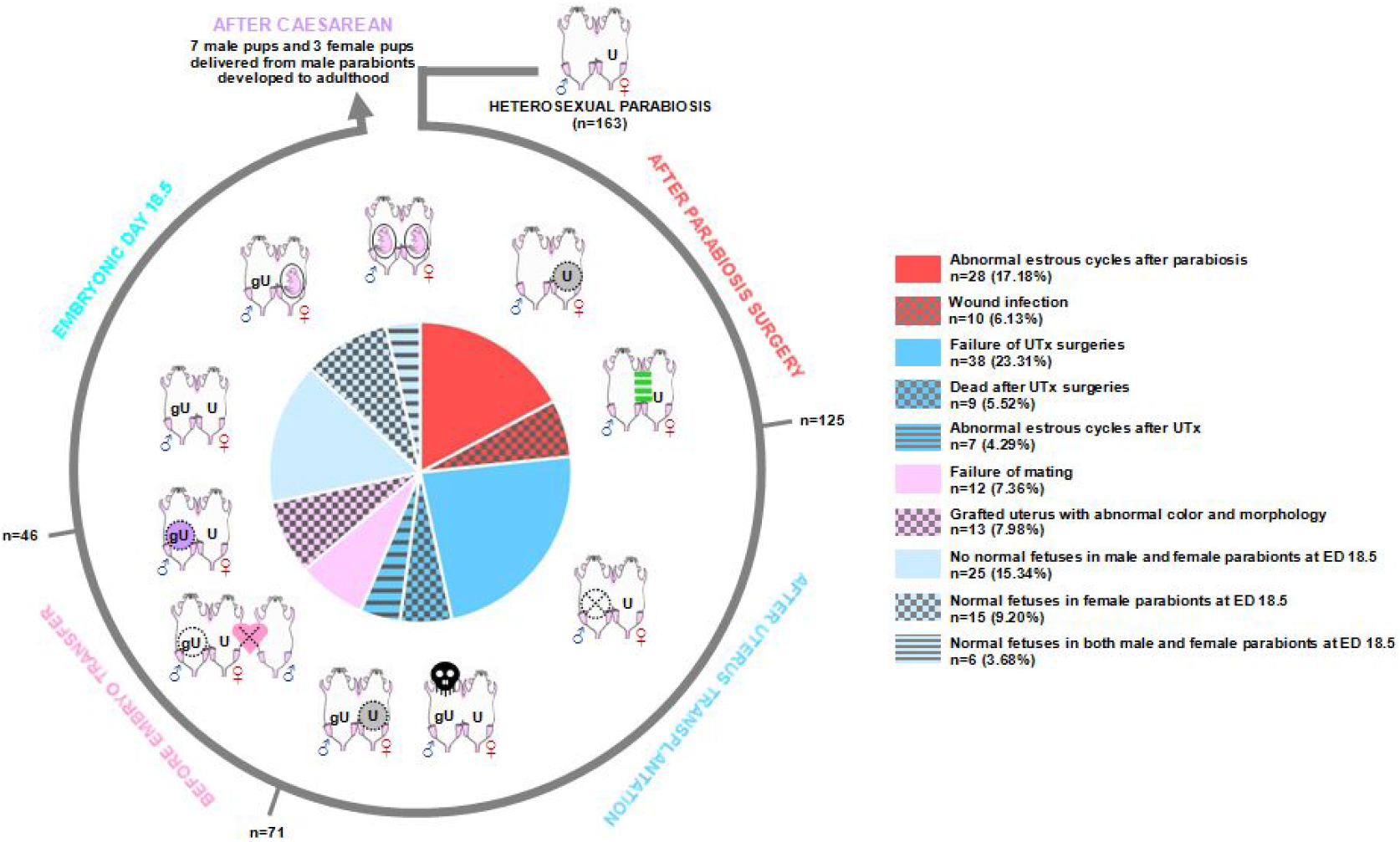
Imaging procedure in our rat model of pregnancy in the male parabiont. In total, 163 heterosexual parabiotic pairs were selected for pregnancy in the male parabiont; only six male parabionts were successfully impregnated (3.68%). In total, 280 blastocyst-stage embryos were transferred into grafted uteruses of male parabionts at ED 4.5; only 27 normal fetuses were observed at ED 18.5 (9.64%). Following caesarean section, 10 pups survived and developed to adulthood (3.57%).

